# Adaptive landscapes unveil the complex evolutionary path to mammalian forelimb function and posture

**DOI:** 10.1101/2024.03.12.584484

**Authors:** Robert J. Brocklehurst, Magdalen Mercado, Kenneth D. Angielczyk, Stephanie E. Pierce

**Affiliations:** Museum of Comparative Zoology and Department of Organismic and Evolutionary Biology, Harvard University, Cambridge, MA 01239, USA; Department of Organismal Biology and Anatomy, University of Chicago, 1027 East 57th Street, Chicago, IL 60637, USA; Negaunee Integrative Research Center, Field Museum of Natural History, 1400 South Lake Shore Drive, Chicago, IL 60605-2496, USA

## Abstract

The ‘sprawling-parasagittal’ postural transition is a key part of mammalian evolution, associated with sweeping reorganization of the postcranial skeleton in mammals compared to their forebears, the non-mammalian synapsids. However, disputes over forelimb function in fossil synapsids render the precise nature of the ‘sprawling-parasagittal’ transition controversial. We shed new light on the origins of mammalian posture, using evolutionary adaptive landscapes to integrate 3D humerus shape and functional performance data across a taxonomically comprehensive sample of fossil synapsids and extant comparators. We find that the earliest pelycosaur-grade synapsids had a unique mode of sprawling, intermediate between extant reptiles and monotremes. Subsequent evolution of synapsid humerus form and function showed little evidence of a direct progression from sprawling pelycosaurs to parasagittal mammals. Instead, posture was evolutionarily labile, and the ecological diversification of successive synapsid radiations was accompanied by variation in humerus morphofunctional traits. Further, synapsids frequently evolve towards parasagittal postures, diverging from the reconstructed optimal evolutionary path; the optimal path only aligns with becoming increasingly mammalian in derived cynodonts. We find the earliest support for habitual parasagittal postures in stem therians, implying that synapsids evolved and radiated with distinct forelimb trait combinations for most of their recorded history.

## Introduction

The evolution of mammals is an iconic transition in the history of life that involved the profound modification of key body systems - feeding^1^, hearing^2^, integument^3^ and metabolic physiology^4^. The exceptionally rich fossil record of mammals and their stem lineage, the non-mammalian synapsids (NMS), documents the assembly of these traits in great detail over some 300 million years^5,6^. Of particular interest is the dramatic reorganization of the ancestral synapsid postcranial musculoskeletal system, including: regionalization of the backbone; simplification of the shoulder girdle; evolution of novel joint types (e.g., ball-and-socket shoulder, trochlear elbow); major restructuring of the limb musculature; and reorientation of the limbs from a horizontal to vertical plane^7–10^. These broad-scale anatomical transformations are intimately associated with a functional shift in limb posture and locomotion, from sprawling pelycosaur-grade synapsids to parasagittal therian mammals.

Although the synapsid ‘sprawling-parasagittal’ transition was a key event in mammalian evolution, its precise nature remains controversial^5,11–14^. Historical studies portrayed synapsid evolution as a stepwise, linear progression towards the therian condition, but often disagreed over when and how major anatomical changes translated into functional or postural change^5,11,12^. These issues primarily arose because previous authors focused on “exemplar” fossil taxa at different key nodes in the synapsid phylogeny and were restricted to qualitative functional interpretations of bony morphology. Recently, more taxonomically comprehensive morphometric work on the synapsid forelimb suggests a pattern of successive evolutionary radiations, with major synapsid groups exploring distinct morphologies and presumed^13,15^. However, to date, few studies on synapsid postcrania have incorporated explicit links between form and function into an analytical framework^16,17^, a crucial step in characterizing the origins of therian-like limb posture and parasagittal locomotion.

Here we use the forelimb, particularly the humerus, as a lens to study postural evolution in synapsids. Forelimb modifications were key to the evolutionary success of synapsids, including mammals^7,8,13–15^, and the humerus provides an important window into forelimb function and posture: it is the primary articulation point between the limb and body, anchors the major muscle groups that drive locomotion, and the arc of motion directly impacts limb movements^10,18–20^. Further, studying isolated humeri allows extensive taxonomic sampling to produce a comprehensive evolutionary viewpoint on the ‘sprawling-parasagittal’ transition. We analyze humerus shape and function across >200 extant and extinct taxa using the integrative analytical framework of evolutionary adaptive landscapes^17,18,21^. This framework links form to functional performance across multiple traits, permitting inference between morphology and higher-level functions such as limb posture or locomotion^18,22^. We extend this synthesis with Pareto optimality analyses^23^ to examine how synapsids transitioned between performance peaks and adaptive optima throughout their evolution and across the ‘sprawling-parasagittal’ postural shift.

Given the differing biomechanical requirements of sprawling vs. parasagittal locomotion^14,24,25^ and the historical perception of a progression towards therian parasagittal posture^5,11,12^, we tested the following hypotheses: 1) Humeri from sprawling and parasagittal taxa occupy different adaptive optima, each maximizing traits most relevant to its specific limb posture and gait; 2) “Pelycosaurs’, the earliest-diverging NMS, share an adaptive optimum with extant sprawling taxa; 3) Increasingly derived NMS taxa shift their adaptive landscapes towards extant therians as they achieve more parasagittal postures; and 4) Evolutionary changes in NMS morphology, function and posture followed an optimal pathway from sprawling to parasagittal. Our data reveal morphological and functional similarities between NMS and extant comparative taxa, but also some key differences. We find that certain performance traits strongly correlate with posture, whereas others relate to different aspects of forelimb function. Recovered patterns show that postural evolution within Synapsida was complex, and the ‘sprawling-parasagittal’ transition was characterized by homoplasy and functional variation within individual synapsid clades, indicative of multiple adaptive radiations of non-mammalian synapsids. Morphofunctional traits consistent with fully parasagittal posture evolved late in the evolutionary lineage of mammals, and so for the majority of synapsid history, the forelimbs were characterized by patterns of variation in form, function and posture that are distinct from what we see in therians today.

## Results

### Humerus shape variation

To capture morphological evolution of the humerus throughout synapsid evolution, we measured humeri from 70 fossil taxa and compared them to 141 extant quadrupedal tetrapods including amphibians, reptiles, and mammals (see Supplementary Table 1 and Supplementary Figure 1). Shape variation was quantified using a novel, homology free pseudo-landmarking approach to generate 3D coordinates along the surface of each bone (see Methods and Supplementary Figure 2). Procrustes aligned landmarks were ordinated using between-groups principal components analysis (bgPCA) to differentiate postural groups (sprawling vs. parasagittal vs. unknown for fossils) and produce a morphospace (Figure 1). Humeri of major extant groups fall in distinct regions of morphospace, but the main axis of shape variation (bgPC1) does not separate them based on posture. Instead, bgPC1 differentiates the relatively gracile humeri of parasagittal therians mammals and sprawling “herptiles” (non-avian reptiles plus amphibians), from the robust humeri of sprawling monotremes (Figure 1A, B). Therians separate from herptiles along the second axis of variation (bgPC2), which reflects differences in the offset between the proximal and distal ends of the humerus, curvature of the humeral shaft and relative width of the proximal vs. distal ends of the humeral epiphyses (Figure 1A, B).

**Figure 1.**
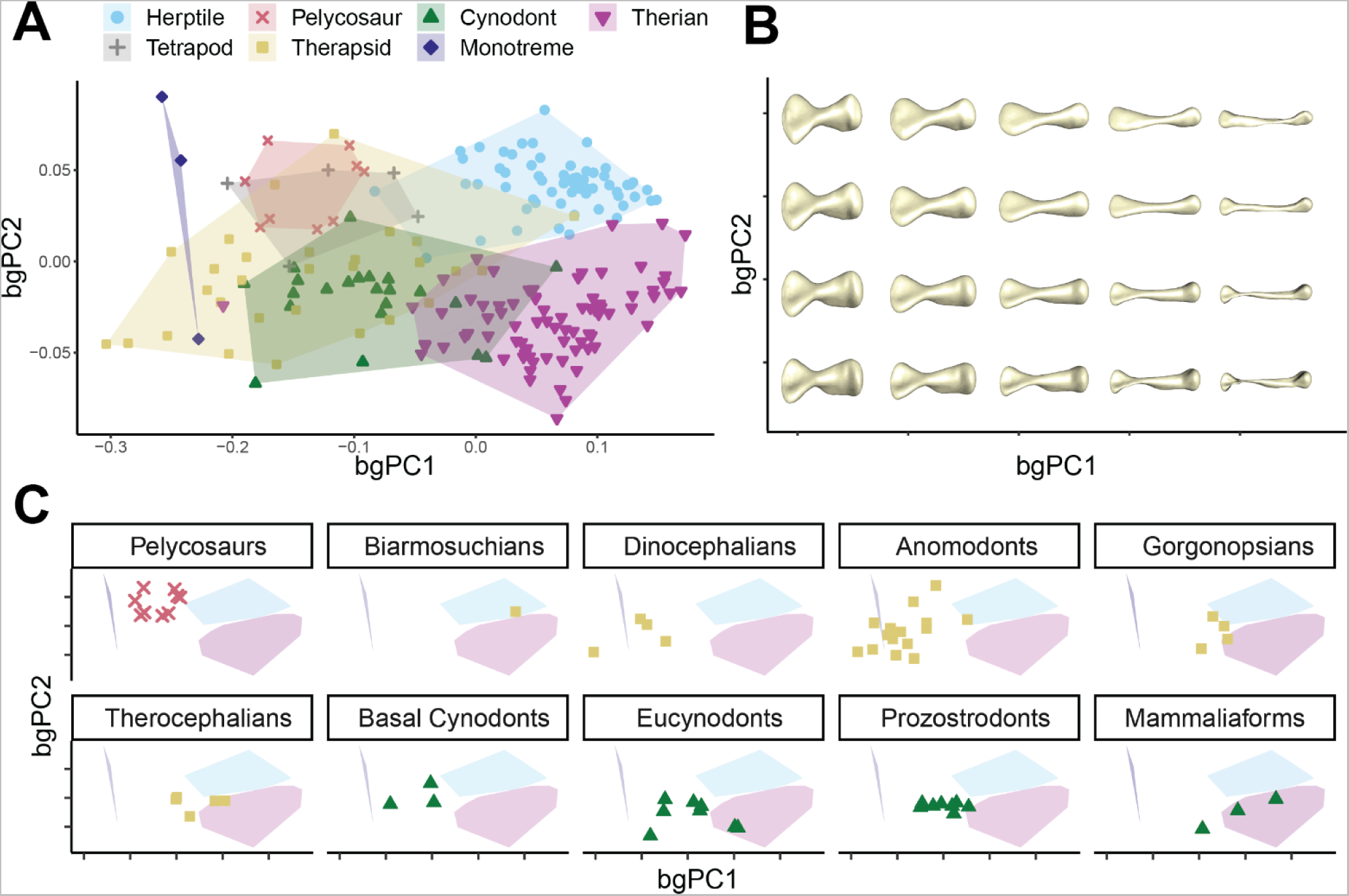
Humerus morphological variation. (A) Humerus between-groups PCA morphospace showing major axes of shape variation. (B) Warps illustrate representative humerus morphologies at each point in morphospace. (C) Subplots show positions of non-mammalian synapsid subgroups, relative to the convex hulls of the three extant groups. See supplementary information for a comparison between PCA and bgPCA results.

Pelycosaurs, the earliest grade of NMS, occupy the region of morphospace between herptiles and monotremes, while overlapping neither, consistent with their “basal” phylogenetic position, and implying a unique sprawling posture^12,26^. Other non-synapsid fossil taxa also plot in this part of morphospace (e.g., *Eryops, Orobates, Seymouria*^19,21^), indicating pelycosaur humeri do not deviate strongly from a general early crown-tetrapod condition. However, more derived grades of NMS – therapsids and cynodonts – diversify into much larger regions of morphospace and exhibit greater variation in humerus shape (Figure 1A, C and Supplementary Figure 3).

In therapsids, this variation is partitioned across different subclades. Anomodonts and dinocephalians generally plot with pelycosaurs or closer to monotremes. The biarmosuchian *Hipposaurus*, our earliest-branching therapsid, plots within herptiles, gorgonopsians plot on the outer edges of the therian and herptile spaces, and therocephalians plot close-to or within the therian cluster (Figure 1C). Basal cynodonts generally occupy the central region of morphospace, as do the two more derived eucynodont subclades, cynognathians and probainognathians, but in both subclades some species independently move into the therian region of morphospace (see Supplementary Figure 3). Some mammaliaforms - *Eozostrodon* and *Megazostrodon*^27^ - fall within therian morphospace, but others - *Borealestes*^28^ - do not (Figure 1C, see Supplementary Figure 3). The two stem therians included here fall well within the range of extant, crown-therian morphologies (Figure 1A, see Supplementary Figure 3).

### Functional Traits, Performance Surfaces and Trait Optimization

Function was quantified by measuring seven osteological proxies - bone length, rotational inertia, torsion, strength, and muscle force and speed leverage - for each humerus in our sample (see Methods). We interpolated the measured functional traits across the morphospace^29^, to produce seven distinct performance surfaces that illustrate the co-variation of humerus shape with each functional trait (Figure 2A). Performance surfaces were combined to produce adaptive landscapes for each species in the dataset using combinatorial optimization, weighting functional traits to maximize each species’ height on their resulting landscape^17,18,21^. Combinations of trait weights that produced optimal landscapes were then compared across taxa and along the synapsid phylogeny (Figure 2B, C).

**Figure 2.**
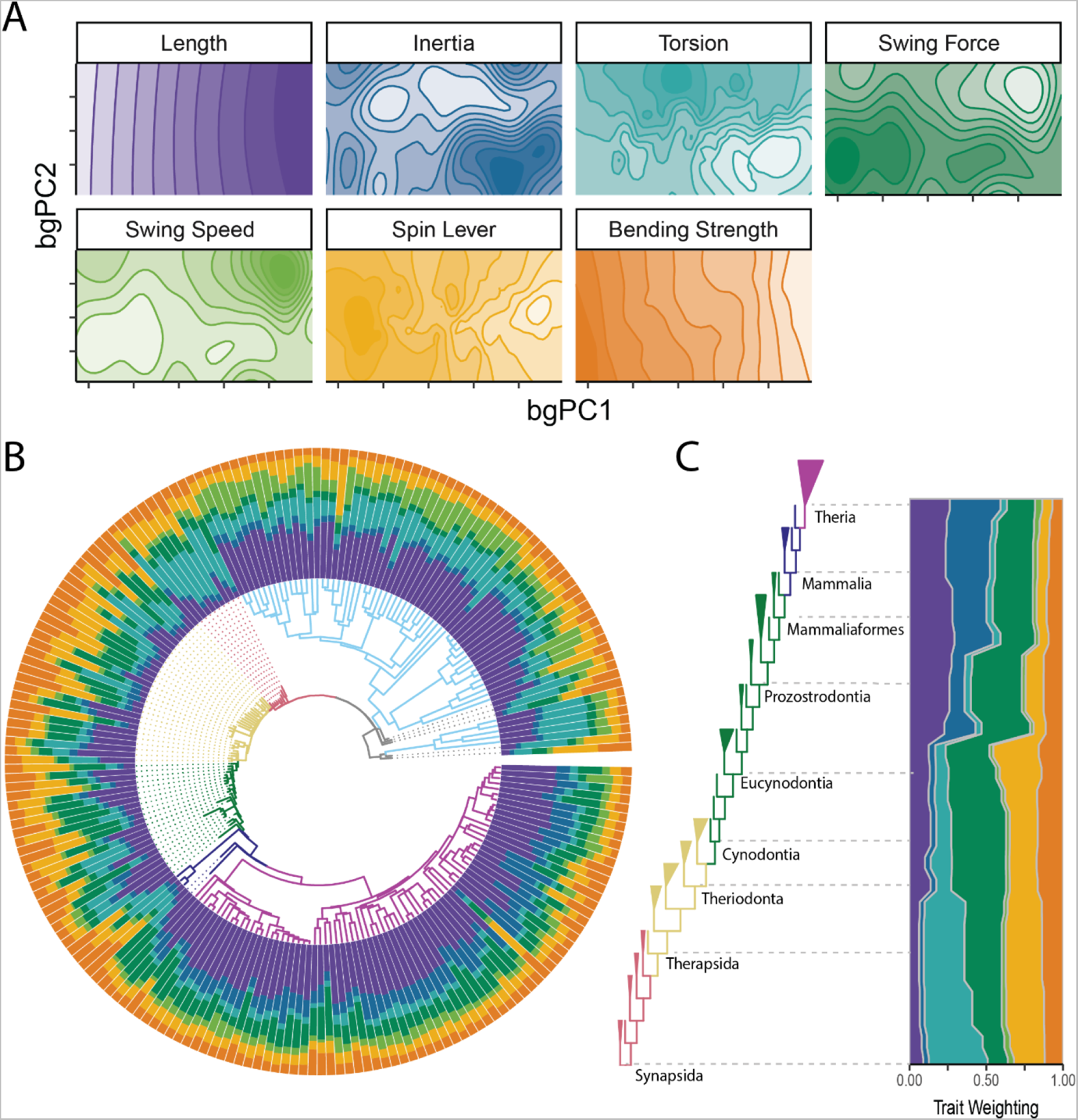
Functional trait variation. (A) Performance surfaces of seven measured functional traits. (B) Trait weightings of adaptive landscapes calculated for individual species plotted onto the phylogeny. (C) Weightings of adaptive landscapes reconstructed at ancestral nodes plotted against a simplified phylogeny of non-mammalian synapsids.

Species-level analysis of trait weights shows considerable variation across, but also within major groups, especially within non-mammalian synapsids (Figure 2B, Supplementary Figure 4). Examining adaptive landscapes based on ancestral-state reconstructions of humerus shape along the backbone of the synapsid phylogeny (Figure 2C) reveals the relative importance of individual traits through synapsid evolution, semi-independent of clade- or taxon-specific patterns. The reconstructed ancestral landscape for the Synapsida node heavily optimizes humeral torsion, as well as ‘spin’ muscle leverage, and this combination remains stable throughout early synapsid evolution. The landscape changes at the base of Theriodontia (gorgonopsians, therocephalians and cynodonts), and throughout Cynodontia, with an increase in optimization for humeral length, ‘swing’ leverage and a decrease in weighting for torsion (Figure 2C). Significant changes to the reconstructed landscapes occur prior to the origins of prozostrodontian cynodonts and Mammaliaformes: increased optimization for humeral length, rotational inertia, and ‘swing’ leverage, with less of an emphasis on humeral torsion and ‘spin’ leverage (Figure 2C).

In addition to individual adaptive landscapes for each specimen, we calculated adaptive landscapes for major taxonomic groups within our dataset; here, the group mean is maximized on the resulting landscape. Extant species optimize different combinations of functional traits (Figures 2, 3, Supplementary Figure 4). Herptiles strongly emphasize humerus length and humeral torsion (Figure 3), traits associated with increasing stride length in sprawling tetrapods^30,31^, as well as high muscle velocity advantage (Figure 3), resulting in fast limb movements and greater muscle working range^14^. Monotremes show high weighting for humeral torsion, ‘spin’ muscle leverage for humeral long-axis rotation, and bending strength (Figure 3), reflective of both their sprawling posture and semi-fossorial lifestyle^14,32–34^. Therians optimize humerus length, rotational inertia, and ‘swing’ muscle leverage for rotating the limb through an arc in a plane (Figure 3). Length and inertia are associated with more efficient locomotion^35,36^, whereas increasing muscle leverage for planar rotation would result in more powerful movements of the limb in the parasagittal plane^14,37^.

**Figure 3.**
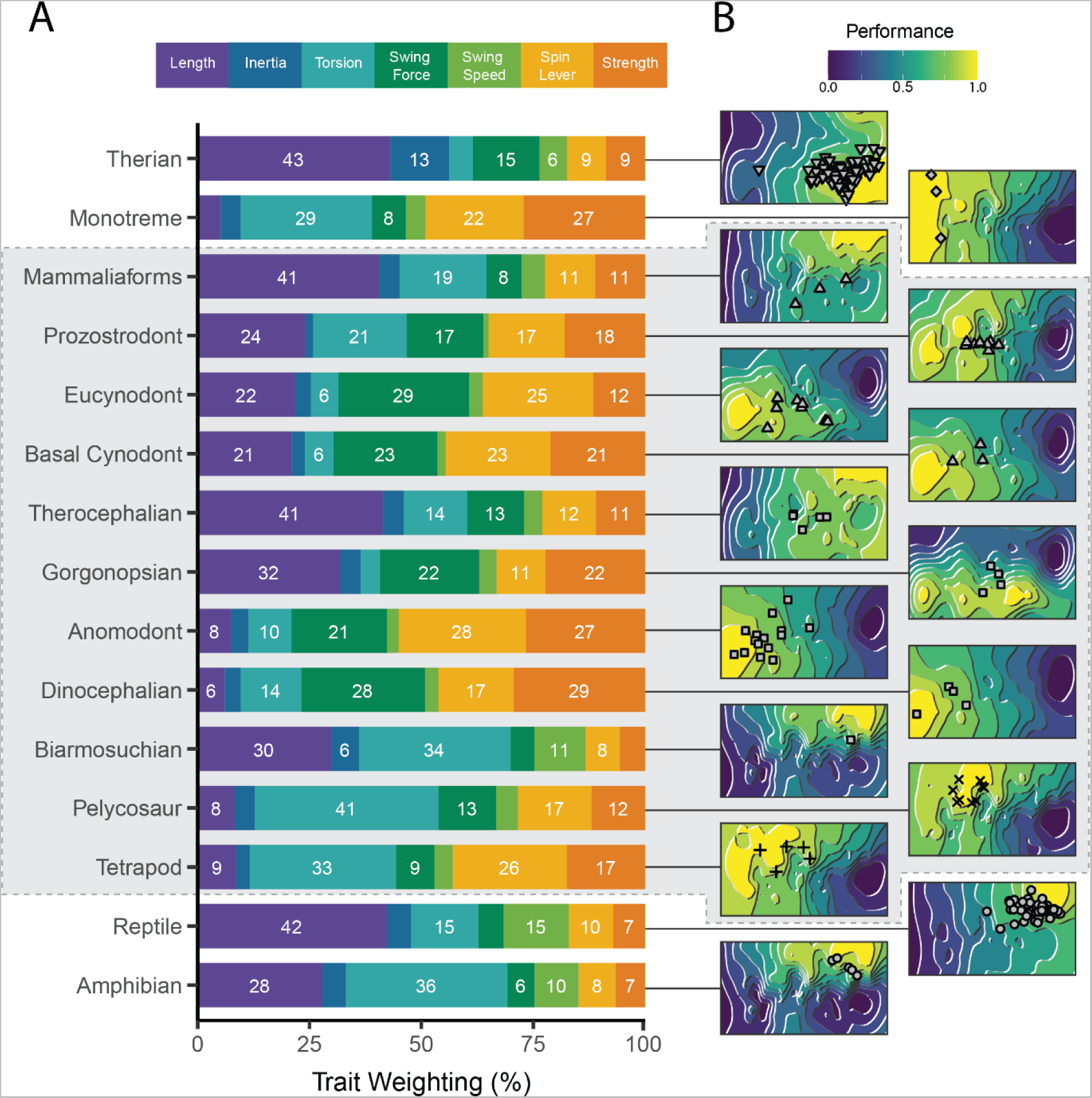
Adaptive landscapes and underlying trait weights. (A) Trait weightings of adaptive landscapes calculated for major extant and fossil groups. (B) The reconstructed adaptive landscapes for each group, showing performance peaks and valleys across morphospace. Fossil taxa are indicated by the grey outline.

Individual NMS vary significantly in how different functional traits are optimized on their landscapes (Figure 2B, Figure 3, Supplementary Figures 4, 5). Pelycosaurs primarily emphasize humeral torsion, consistent with their reconstructed sprawling posture^38,39^. Other traits – spin leverage, swing force, and humeral strength – also contribute to the pelycosaur landscape with varying importance across taxa (Figure 3, Supplementary Figures 5). Therapsids differ in functional trait optimization across subclades (Figures 2B, 3, Supplementary Figures 4, 5). The biarmosuchian *Hipposaurus* emphasizes torsion and humeral length, similar to herptiles. Dinocephalians and anomodonts predominantly optimize strength and muscle leverage. Gorgonopsians and therocephalians optimize humeral length, but whereas gorgonopsians emphasize swing torque and strength, therocephalians put more importance on humeral torsion (Figure 3, Supplementary Figure 5). Cynodonts show substantial variability in trait optimization, but generally emphasize either humerus length, or a combination of strength and muscle leverage (Figures 2B, 3, Supplementary Figure 4, 5). Most cynodonts do not optimize humeral torsion, but there are some exceptions including among derived prozostrodonts and mammaliaforms (Supplementary Figure 4, 5). This contrasts with therian trait optimization, as well as that reconstructed along the synapsid backbone – the importance of torsion in these taxa may represent independent instances of trait diversification.

### Transitions from Sprawling to Parasagittal Postures

To further investigate patterns of humeral transformation towards (or away from) the therian condition during the evolution of Synapsida, we created a transitional ‘sprawling-parasagittal’ landscape. We first created a composite “sprawling” landscape by combining the independently calculated landscapes for different sprawling groups (Figure 4A and see Methods). Then, for each point in morphospace, we subtracted the height on the “sprawling” landscape from the height on the therian (i.e., “parasagittal”) landscape (Figure 4B) to create the transitional landscape (Figure 4C). Transitional landscape scores indicate the relative performance of different humerus morphotypes for sprawling vs. parasagittal limb functions, given the many-to-one mapping that exists between humeral traits and posture^19,32,40^. We plotted our taxa on this new ‘sprawling-parasagittal’ landscape (Figure 4C, D), determined their adaptive score (height), mapped these scores onto a phylogeny and reconstructed ancestral states using maximum likelihood (Figure 5). Evolutionary shifts in transitional landscape score within Synapsida were identified using the SURFACE algorithm (see Methods).

**Figure 4.**
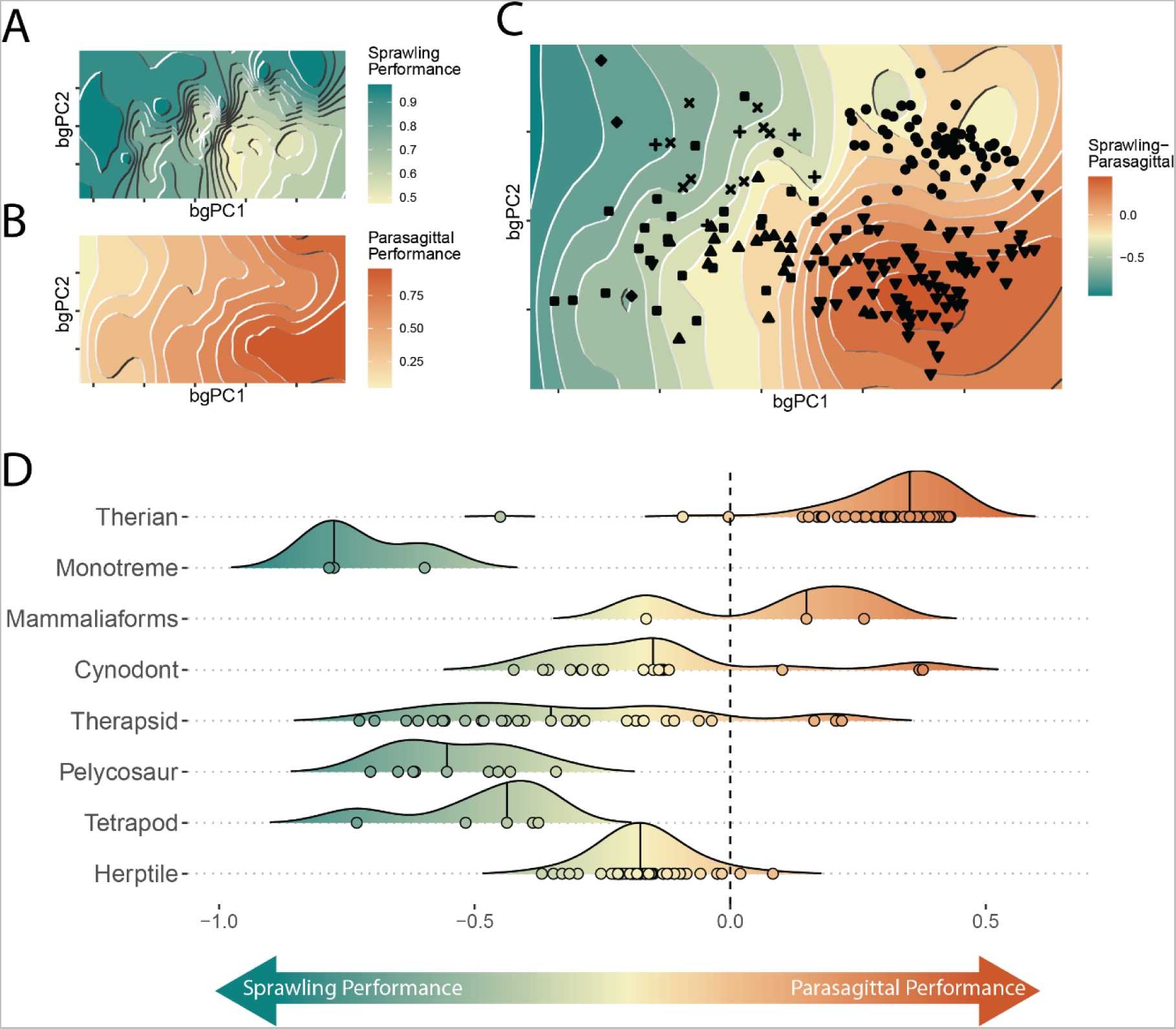
Transitional ‘sprawling-parasagittal’ landscape. (A) Composite sprawling landscape (reptiles, salamanders, monotremes and non-synapsid fossils), and (B) parasagittal (therian) landscape, (C) overlaid to create a transitional ‘sprawling-parasagittal’ landscape. (D) Scores on the transitional ‘sprawling-parasagittal’ landscape plotted for major extant and fossil groups.

**Figure 5.**
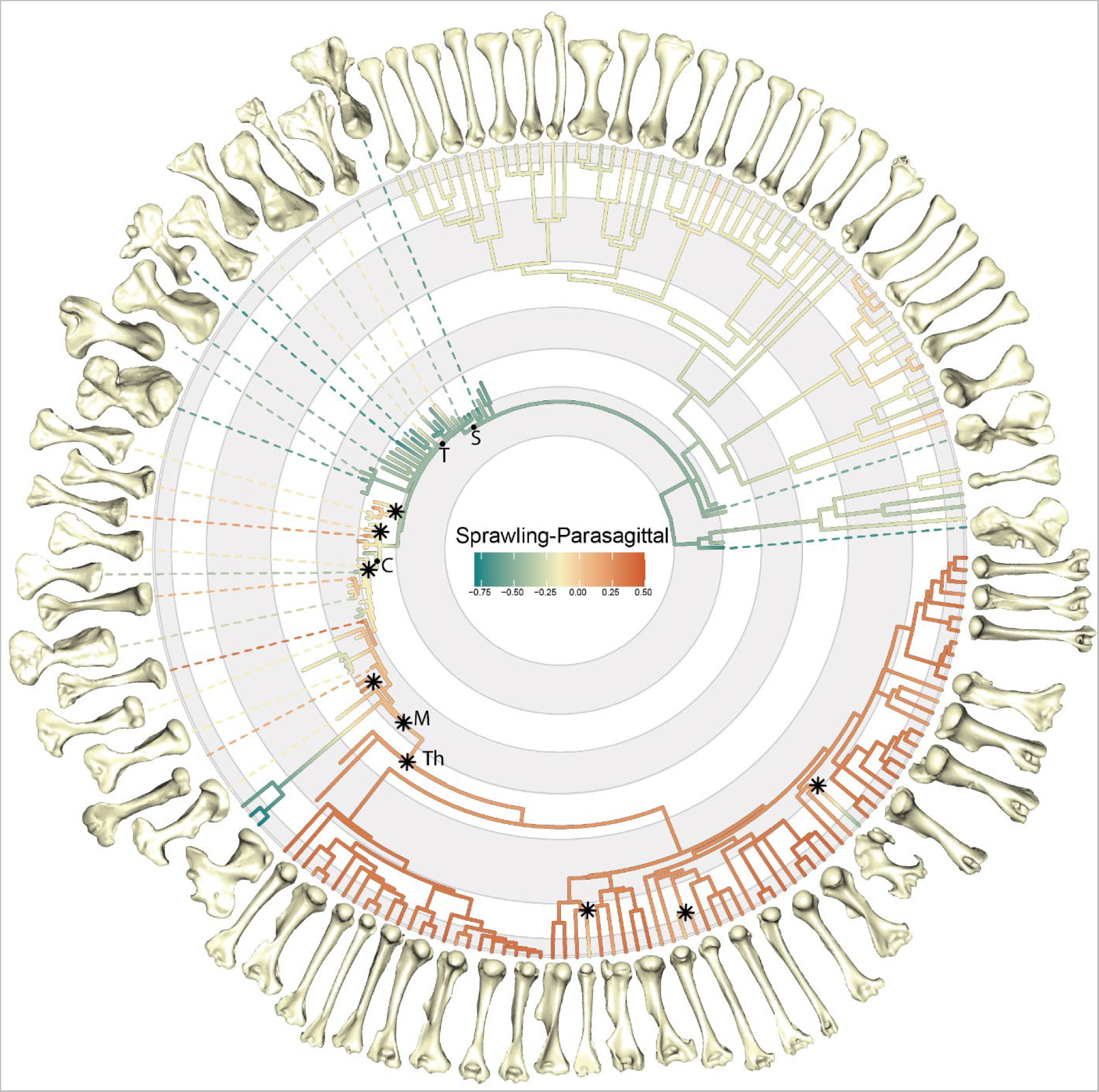
Evolution of sprawling and parasagittal posture. Scores on the transitional ‘sprawling-parasagittal’ landscape are plotted on the phylogeny of all specimens, illustrated with select humeri. Stars indicate shifts in evolutionary regime for Synapsida identified by SURFACE (see Methods). Nodes labelled with letters; S, Synapsida, T, Therapsida; C, Cynodontia; M, Mammalia; Th, Theria

As expected, therians had the highest scores and monotremes had the lowest, indicating trait combinations most and least associated with parasagittal postures respectively (Figure 4C, D). Herptiles were intermediate between monotremes and therians, due to sharing morphofunctional traits with both groups (humeral torsion with monotremes, and long humeri with therians), but still had negative scores indicating greater association with sprawling locomotion (Figure 4D). Thus, the transitional landscape helps to conceptualize posture as a continuous variable; sprawling and parasagittal represent extremes, but we also show the great variation present within sprawlers and multiple paths between different postural groups^19,40,41^. Ancestral state reconstruction recovers the ancestor of crown-mammals as closer to therians than monotremes (Figure 5). Consequently, extant sprawling monotremes do not retain a plesiomorphic state for Synapsida but have rather converged on a suite of morphofunctional characters that resemble ancestral, sprawling synapsids. This convergence may have been facilitated by the incomplete acquisition of therian characters in early mammals and their close relatives (e.g., retention of humeral torsion in taxa such as *Brasilodon*, *Morganucodon*, and *Gobiconodon*).

Our analyses reconstruct ancestral humeri at the base of Synapsida with traits consistent with sprawling postures, which persist throughout the pelycosaur and early therapsid parts of the tree (Figure 4D, 5). Pelycosaurs, dinocephalians and most anomodonts have humeri with very negative scores on the transitional landscape, providing strong evidence for sprawled postures (Figure 5, Supplementary Figure 6). Although our analysis indicates a sprawled ancestor for Therapsida with monotreme-like traits, several early therapsids show less negative scores (e.g., *Hipposaurus, Tiarajudens),* more in-line with modern reptile values, possibly indicative of differences in locomotion (e.g., faster limb movements and different kinematics) (Figure 4D, 5, Supplementary Figure 6).

We see independent shifts towards intermediate postural scores in gorgonopsians and therocephalians (Figure 5, Supplementary Figure 7). Although these groups have humeri with distinctly higher transitional landscape scores than earlier synapsids, their scores are generally still negative, like modern reptiles. Another shift on the transitional landscape occurs at the origin of Cynodontia, with most taxa possessing intermediate postural scores (Figure 4D, 5, Supplementary Figures 6, 7). However, there is considerable heterogeneity within cynodonts and later mammaliaforms, with taxa evolving humeri with both more negative (e.g., *Chiniquodon*, *Exaeretodon*) and more positive scores (e.g., *Massetognathus*, *Probainognathus*) (Figure 5, Supplementary Figure 6). By contrast, within crown mammals, the two stem therians (*Adalatherium* and *Gobiconodon*) both scored positively on the transitional landscape (Figure 5).

### Performance Trade-offs and Optimal Evolutionary Pathways

To determine whether synapsids followed “optimal” paths through morphospace over the course of their evolution as they transitioned from one adaptive optimum to another, we calculated Pareto optimality landscapes^23,42^. Trade-offs necessarily occur between high fitness on different landscapes due to differences in the underlying functional traits being optimized. Pareto optimality finds points in morphospace whose height on one landscape is maximized, given their height on another landscape^23,43^. By combining adaptive landscapes reconstructed for major nodes in the synapsid phylogeny, we determined if taxa evolve optimally through morphospace from an ancestral peak to that of the next more derived clade (see Methods). Deviation from high Pareto optimality implies exploration of novel morphologies and functions, and either the presence of new functional drivers distinct from those along the synapsid backbone or the weakening of functional constraints and lowering of restrictions on limb evolution over time^23^.

Contrary to expectations that it would follow an optimal path towards a more therian morphology, the backbone of the synapsid phylogeny shows some fluctuations in optimality over time, based on reconstructed landscapes for major nodes with individual clades branching off to explore both more and less optimal regions of morphospace (Figure 6, Supplementary Figure 8). Pelycosaurs undergo a small radiation but are mostly restricted to regions of high optimality that connect their adaptive peak with that for the reconstructed therapsid ancestor (Figure 6). In therapsids, several clades – biarmosuchians, gorgonopsians and therocephalians – explore parts of morphospace with lower optimality on the landscape connecting Therapsida with Cynodontia (Figure 6, Supplementary Figure 8). Dinocephalians and anomodonts both exhibit greater Pareto optimality on this landscape, clustering around the inferred Therapsida optimum (Figure 6, Supplementary Figure 8). Basal cynodonts and eucynodonts generally occupy regions of lower Pareto optimality, but certain taxa evolve towards distinct peaks on the cynodont-prozostrodont landscape, with some species evolving towards the reconstructed ancestral cynodont optimum, associated with more robust humeri, and others evolving towards the prozostrodont optimum, associated with more gracile humeri (Figure 6). Finally, prozostrodont cynodonts and mammaliaforms also occupy sub-optimal regions, although there does seem to be a shift in morphospace occupation towards the optimum reconstructed for ancestral therians (Figure 6, Supplementary Figure 8).

**Figure 6.**
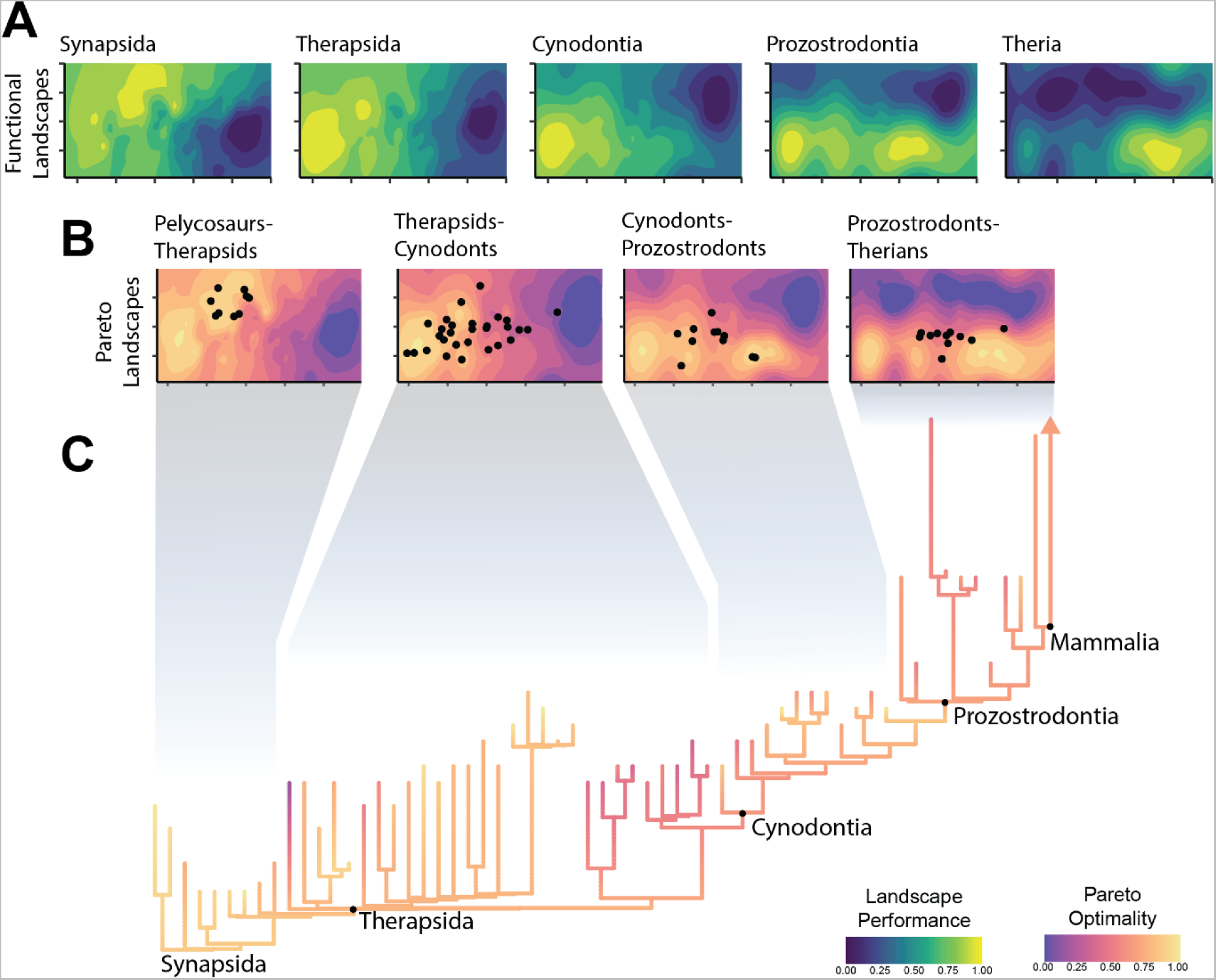
Pareto optimality across synapsid evolution. (A) Functional adaptive landscapes reconstructed for major ancestral nodes within synapsid evolution and (B) the Pareto landscapes created by combining these landscapes together. Pareto optimality for a group of taxa is defined based on their ancestral peak, and the peak of the next major node. (C) Pareto optimality plotted on the non-mammalian synapsid phylogeny.

## Discussion

The evolution of parasagittal posture in mammals and their ancestors has been studied for over a century, but previous efforts to understand changes to the limbs and locomotion have historically lacked taxonomic scope and an appropriate analytical framework^5,11,12^. Here, we used evolutionary adaptive landscapes^17,21^ to predict relationships between humerus morphology, function, and posture to develop a predictive framework to illuminate the ‘sprawling-parasagittal’ transition in synapsids. Based on differences in locomotor biomechanics, we expected the humeri of extant taxa to group together in morphospace based on posture (Hypothesis 1). Although parasagittal therians group separately from other taxa, sprawling reptiles and amphibians (“herptiles”) do not group with sprawling monotremes. Together with therians, herptiles share long, gracile humeri with strong emphasis on humerus length (Figures 1 and 3, Supplementary Figure 4), features that arose convergently in these groups despite their contrasting habitual postures. Longer humeri are likely advantageous for general terrestrial locomotion regardless of posture, although the precise selective advantages may differ (e.g., speed vs. efficiency)^30,35,36^. In contrast to therians, herptiles and monotremes both emphasize humeral torsion, supporting torsion as a strong indicator of sprawling forelimb posture^31,44^ (Figures 2-4). However, monotremes and herptiles optimize torsion to differing degrees and in distinct combinations with other functional traits, reflecting wide variation in sprawling locomotor kinematics (e.g., powerful long-axis rotation vs. fast limb retraction)^19,32,33^. Therians are unique in emphasizing rotational inertia (Figures 2, 3), indicating this trait as a likely signal of more parasagittal postures. These similarities and differences across diverse extant animals provide interpretive power when reconstructing forelimb function and postural evolution in synapsids.

The earliest-diverging NMS, the pelycosaurs, are traditionally reconstructed with a sprawling posture^38^, and thus we hypothesized morphological and functional similarities with extant sprawlers (Hypothesis 2). Yet, our analysis shows pelycosaurs are morphologically distinct from all extant sprawling groups, and instead overlap with extinct non-synapsid tetrapods, indicating that pelycosaur humeri had not diverged from the plesiomorphic crown tetrapod condition (Figure 1). Functionally, pelycosaurs show high optimization for torsion, indicative of sprawling posture, but they optimize this trait to a more extreme degree than either reptiles or monotremes (Figure 3, Supplementary Figure 4). Pelycosaurs also emphasize a combination of both swing and spin muscle leverage, which aligns with previous work demonstrating coupled rotations at the screw-shaped glenoid joint during a stride^12,26^. Taken together, the emphasis on torsion and muscle leverage shows pelycosaurs used slow, forceful limb movements, combining humerus long-axis rotation and retraction. This combination contrasts with reptiles, which are adapted for speed^14^, and monotremes, which use almost exclusively long-axis rotation^32,33^. Our analyses thus indicate pelycosaurs possessed distinct humerus morphologies and combinations of functional traits not represented by modern taxa^17^, and it was from this unique starting point that mammalian posture ultimately evolved.

Based on historical interpretations of the synapsid fossil record^5,11,12^, we expected a trend towards more therian-like morphologies and functional traits in more derived NMS, reflecting more parasagittal postures (Hypothesis 3). Although this pattern was broadly supported, we also found significant variations on this theme (Figures 1-3). Our transitional ‘sprawling-parasagittal’ landscape strongly supports monotreme-like sprawling postures in dinocephalian and anomodont therapsids (Figures 4 and 5), with these clades showing increased optimization for strength and muscle leverage at the expense of humerus length (Figures 2 and 3, Supplementary Figures 4, 5). This combination of functional traits is potentially related to fossorial behaviors in smaller dicynodonts^45–47^, as seen in modern monotremes and talpid moles^32,33,48^, and the acquisition of larger body sizes in both groups^49,50^. Similar ecomorphological convergence in the forelimb has been previously noted between fossorial and large-bodied mammals^13^. We argue that the co-occurrence of large body sizes and strong humeri in dinocephalians and anomodonts is indicative of non-parasagittal postures in these groups. Modern large-bodied therians can compensate for size-related bending stresses by changing limb posture, from more crouched to more erect^24^. Sprawling taxa, on the other hand, must accommodate size-related stresses by increasing limb bone robusticity, resulting in “overbuilt” limb bones^25,51^.

Other therapsid groups had diverging scores on the ‘sprawling-parasagittal’ transitional landscape. Biarmosuchians, the earliest branching therapsids and represented here by *Hipposaurus*, experienced a shift on the transitional landscape away from monotremes and towards reptiles (Figures 4 and 5, Supplementary Figures 6, 7). Specifically, *Hipposaurus* overlaps with reptiles in morphospace and optimizes humeral torsion, length and ‘swing’ speed (Figures 1 and 3), indicating a reptile-like sprawling posture and kinematics^19^. Although this combination suggests adaptations for fast limb movements, it may not be reflective of the ancestral therapsid condition, due to a persistent ghost-lineage following their divergence from pelycosaurs^52^. Later theriodont therapsids, the gorgonopsians and therocephalians, experienced similar but less extreme shifts on the transitional landscape, sitting on the outskirts of the reptile and therian regions of morphospace (Figures 4, 5, Supplementary Figures 6, 7). Therocephalians share similar functional traits to *Hipposaurus*, but gorgonopsians combine humerus length with swing force and strength. These morphofunctional traits are indicative of more active, predatory lifestyles in the three clades^53–56^, but further suggests that gorgonopsians may have engaged in unique behaviors with their forelimbs (e.g., grappling large prey similar to more recent carnivores^57^). Therocephalians and gorgonopsians both have average clade-wise postural scores converging on reptiles, but several taxa (*Gorgonops*, *Olivierosuchus*) have higher scores closer to therians (Supplementary Figure 6). This convergence may reflect more parasagittal postures in these taxa, but the range of forelimb poses would have been constrained by the therapsid shoulder girdle, with its caudolaterally facing glenoid^5^.

Following theriodonts, further morphofunctional transformations to the forelimb associated with increasingly parasagittal postures have been proposed for cynodonts^5,12^. Cynodonts do shift closer to therians on the transitional landscape, but this movement is accompanied by considerable variation (Figures 1, 4, Supplementary Figure 6) and evolutionary heterogeneity (Figure 5, Supplementary Figure 7). Whereas postural trait variation in therapsids is phylogenetically structured by clade, in cynodonts it is more widespread across the phylogeny, implying enhanced evolutionary lability in posture and forelimb use. For functional traits, cynodonts generally optimize either humerus length, or a combination of strength and muscle leverage (Figure 2 and 3, Supplementary Figures 4, 5), possibly reflecting adaptations to distinct, specialized ecologies (e.g., digging^58^). Most cynodonts are not optimized for humeral torsion, but those that are (e.g., *Lumkuiia, Riograndia, Brasilodon*) have been previously reconstructed with sprawling postures^59–61^. Likewise, lower torsion might indicate less sprawled postures in other taxa^62^. Postural scores in cynodonts are mainly in-line with modern reptiles, but some taxa achieve higher, more therian postural scores (*Massetognathus*, *Probainognathus*) (Figure 5, Supplementary Figure 6). Given the disparity in cynodont forelimb functional and postural traits, it is likely that different species employed different postures, but were still constrained by the shoulder girdle to non-parasagittal (i.e., sprawling to semi-sprawling) limb poses^12,62–64^. Cynodont forelimb disparity is further explored by mammaliaforms, which possess disparate postural scores and functional traits: humerus length in *Megazostrodon*; length and torsion in *Morganucodon*; and length, spin leverage and strength in *Borealestes* (Figure 2, Supplementary Figure 5). The functional, and likely postural, diversity of mammaliaform humeri reflects the great ecomorphological disparity present in this group^65^.

Our comprehensive survey of non-mammalian synapsids found evidence of therian traits evolving multiple times, but limited support for fully parasagittal postures. Previous hypotheses place the origin of parasagittal posture in crown Mammalia^11^ or even Theria^12^. Our two stem therians (*Adalatherium* and *Gobiconodon*) both plot with crown therians on the transitional landscape (Figures 1, 4). However, when we examine humerus traits, we find *Gobiconodon* strongly optimizes humeral torsion, which would preclude it from adopting a habitually parasagittal posture^66^ (Figure 2, Supplementary Figure 5). *Adalatherium,* on the other hand, possesses a functionally therian humerus, optimizing length, swing leverage and, to a lesser extent, rotational inertia. Further support for therian, parasagittal posture in *Adalatherium* comes from other aspects of the forelimb skeleton: the scapula has a ventrally facing glenoid facet, and the coracoid portion of the glenoid fossa faces caudo-ventrally not laterally^67^. A similar condition in multituberculates, in combination with low humeral torsion, has been interpreted as evidence these taxa were also parasagittal^68,69^. Therefore, our results support the origins of fully parasagittal posture within stem therians^12^, making it a late innovation in the grand scheme of synapsid evolution. Additionally, due to uncertainty in the phylogenetic arrangement of gondwanatherians (e.g., *Adalatherium*), eutriconodonts (e.g., *Gobiconodon*) and multituberculates^70,71^, it is possible that adaptations for more parasagittal postures arose multiple times independently along the therian stem.

From evolutionary theory, we predicted that throughout their complex evolutionary journey, NMS were following an optimal pathway through morphospace (Hypothesis 4). Following prior work on synapsid evolution^5,11,12^, we assumed that this optimal pathway would correspond to the acquisition of increasingly therian traits and postures. Instead, our Pareto optimality analyses clearly illustrate synapsid evolution as a series of successive adaptive radiations (Figure 6, Supplementary Figure 8), with taxa repeatedly evolving away from the reconstructed optimal pathway. Pelycosaurs represent the first radiation of synapsids, but they do not diversify the forelimbs to the same extent as later synapsid grades^15^. The second radiation at the origin of Therapsida corresponds to a proposed morphofunctional shift in synapsid evolution based on important morphological transformations like the change in the scapular glenoid from a screw-shape to a more mobile hemi-sellar shape^5,11^. The removal of morphofunctional constraints on the forelimb potentially facilitated therapsid diversification, as they explored novel regions of morphospace^15^ and experimented with novel forelimb functions^46,56,72^ (Figures 1-3, Supplementary Figure 5). Some clades diversified in areas of high Pareto optimality (e.g., anomodonts), but biarmosuchians and theriodont therapsids explored Pareto suboptimal regions of morphospace and combinations of functional traits closer to those seen in therians and reptiles. This variability gives Therapsids a lower average Pareto optimality than pelycosaurs, providing strong supporting evidence for reduced evolutionary constraints^23^. Recovering theriodont therapsids as Pareto suboptimal also demonstrates that becoming increasingly therian was not the ‘optimal’ evolutionary pathway for synapsids until much later in their history.

Contrary to therapsids, cynodonts and mammaliaforms evolve on a multi-peak Pareto landscape, with optima corresponding to more robust and gracile humeri associated with the ancestral Cynodontia and derived Prozostrodontia landscapes, respectively (Figure 6, Supplementary Figure 8). However, only a few taxa occupy either of these optimal regions, with most cynodonts and mammaliaforms occupying Pareto suboptimal regions between the peaks. We propose that the two peaks, representing distinct humerus morphologies and functional traits, constitute two ends of an ecological spectrum (e.g., fossorial to scansorial^59,60^), with more generalist taxa falling in the center due to conflicting selective pressures^73^. The gracile peak on the prozostrodont Pareto landscape is closer to the therian optimum and forms a Pareto optimal evolutionary pathway towards Theria. However, this is not the only optimal path on this landscape, as moving back towards the ancestral cynodont peak is also Pareto optimal. These multiple optima may have facilitated the disparate evolution of mammaliaforms^65^, and the opposing evolutionary trajectories of monotremes and therians. Monotremes convergently re-evolve towards the robust, ancestral cynodont optimum, and therians realize a new set of humerus morphologies and functions. Accompanied by modifications to other aspects of the therian forelimb skeleton along the therian stem^67^ – a mobile scapula and ventrally facing glenoid – this new morphology presumably removed the final barriers to achieving a habitually parasagittal forelimb posture^14^.

## Conclusion

The evolution of parasagittal posture in Synapsida involved a fundamental reorganization of the musculoskeletal system and expansion of forelimb functional disparity, culminating in extant Theria. Our comprehensive examination of the ‘sprawling-parasagittal’ transition uses the framework of evolutionary adaptive landscapes to quantify differences in humerus morphology and functional traits between sprawling and parasagittal taxa and reconstructs the evolution of form and function in non-mammalian synapsids. Our results indicate that although synapsids were ancestrally sprawling, they were both morphologically and functionally distinct from extant sprawling taxa. From this unique sprawling ancestral condition, we find that synapsids humerus morphofunctional evolution did not follow a single, optimum path to reach the parasagittal condition of modern therians. Some synapsid taxa stayed close to reconstructed group optima, but others diversified through morphospace. More gracile humeri with unique combinations of reptilian and therian traits evolved independently in biarmosuchian, gorgonopsian and therocephalian therapsids, before an extended period of functional experimentation and variation throughout non-mammalian cynodont evolution. We reject previous hypotheses on the synapsid ‘sprawling-parasagittal’ transition as a series of discrete postural shifts, and instead support the view of synapsid evolution as a series of successive radiations, with major clades exhibiting considerable functional (and postural) variation. Our data on humerus morphology and functional trait evolution suggest that parasagittal posture evolved late, within stem therians. More sophisticated biomechanical modelling is required to resolve whole limb posture in specific taxa, but the present study, with its large sample of both fossil synapsids and extant comparators, provides a far more holistic picture of the ‘sprawling-parasagittal’ transition, and the assembly of the therian forelimb, than previous works considering only small numbers of exemplar taxa.

## Methods

### Sample and Phylogeny

Humeri from a range of fossil synapsids (9 pelycosaur, 32 therapsid, 22 cynodont and 2 stem therian specimens, representing at least 58 distinct taxa), were compared to a diverse array of extant taxa (5 salamanders, 56 reptiles, 3 monotremes and 77 therian mammals, Table S1). With respect to understanding the evolution of non-mammalian synapsids, monotremes possess several functional and morphological similarities to fossil taxa^13,32,74^, and extant sprawling reptiles have long been used as analogues for non-mammalian synapsids (NMS) and other fossil amniotes^11,12,19^. Our extant sample covers a great diversity in body size and ecology, but all engage in quadrupedal terrestrial locomotion; volant and fully aquatic taxa were excluded because these derived specializations are not relevant to the ‘sprawling-parasagittal’ transition. We also included a small sample of generalized early fossil amniotes and temnospondyls (n=5) to assist with establishing character polarity for the earliest NMS taxa. All information regarding data provenance, acquisition method, and processing to derive a 3D surface mesh is listed in Table S1.

Relationships and branch lengths among extant species are based on consensus phylogenies downloaded from timetree.org^75^, constructed based on comparison of all published phylogenies of the taxa of interest. Relationships of fossils (including fossil crown-mammals) are based on a combination of two existing large-scale phylogenies of living and extinct synapsids^76,77^, which provide a basic backbone phylogeny of all non-mammalian synapsid fossils. Specimens missing from these two phylogenies were assigned the position of their closest relative present in the trees. The fossil supertree was time calibrated with occurrence data downloaded from the Paleobiology Database (palaeobiodb.org), using the function *timePaleoPhy* from the *paleotree* package in R. The final phylogeny was then constructed by grafting the phylogenies of extant taxa onto the appropriate places in the fossil tree (e.g., salamanders as sister to *Eryops*, reptiles as sister to *Captorhinus* etc.), and pruning the tree to only include sampled fossil taxa.

### Geometric morphometrics and Morphospace

Humerus shape was quantified using a novel semi-automated approach for slice-based landmarking (Supplementary Figure 2). Three-dimensional surface meshes of each humerus were manually aligned to a global coordinate system using a standard protocol in Materialise 3-Matic (v14, Materialise, Leuven, Belgium). The long-axis of the humerus was aligned to the global Z-axis, and the bone was then rotated about its long-axis until the distal medial-lateral epicondyles were horizontal, parallel with the global X axis, and the extensor surface of the humerus (i.e., the face of the humerus to which the elbow extensor muscles attach), faced upwards (see supplemental material). The final sample consisted of only left humeri; any right humeri were mirrored. Once all humeri were consistently oriented, the 3D meshes were converted to ‘.ply’ format and imported into R for landmarking.

Meshes were landmarked with a slice-based approach, using custom code based on the *Morphomap* R package^78^ (Supplementary Figure 2). Each humerus was ‘sliced’ at 21 points evenly spaced along its length, by drawing a plane normal to the long axis that intersected with the bone mesh. In each of the 21 slices, the intersection between the plane and the mesh defined the bone’s outer contour, and equidistant landmarks (n=24) were placed around the contour’s length. This produced a final set of 504 landmarks for each humerus, and these final landmark sets were subjected to general Procrustes alignment in the R package *geomorph*^79^.

Data were ordinated using standard principal components analysis (PCA), which produced generally good separation between taxonomic and postural groups (Supplementary Figure 9). However, to create clearer separation and a more structured morphospace, we also ordinated the data using a between-groups principal components analysis (bgPCA) using the R package *Morpho*^80^. This produced a morphospace that emphasized differences between *a priori* postural groups (sprawling vs parasagittal vs ‘unknown’ for fossil taxa), and the bgPC axes are highly correlated with measured functional traits (Figure 2). Comparison with the traditional PCA shows a similar overall distribution of groups in morphospace, and that our bgPCA did not create ‘false groups’^81^. Therefore, we proceeded with the bgPCA morphospace in subsequent analyses.

### Measuring Functional Traits

#### Functional Humerus Length

Humerus length is an important component of locomotor performance (e.g., speed), as it is a major contributor to overall limb length and longer limbs produce longer strides and greater overall displacement^30^. Although there is considerable ecological variability in humeral length^7^, we predicted that humeral length would increase throughout synapsid evolution. Functional humerus length – the length between the proximal and distal articular surfaces – was measured in 3-Matic as the distance from the proximal-most part of the humeral head, and the distal-most part of the radio-ulnar condyles (Supplementary Figure 10). After normalizing length to humeral centroid size, this differentiated long, gracile humeri from shorter, more robust humeri.

#### Rotational Inertia

Rotational inertia determines how much torque is needed to generate angular acceleration about a rotational axis and is proportional to how much effort it takes to move the limb during locomotion. Rotational inertia is proportional to the mass of the object (m), and the square of the rotational distance, or the distance from the center of mass to the center of rotation (r^2^). Although we could not measure humerus mass directly, we used volume as a proxy. We measured the volume and position of the center of mass of each humerus model in 3-Matic. The center of rotation was determined by fitting primitives (spheres, cylinders, or convex hulls) to the articular surface of the humeral head. We then measured the distance between this point and the center of mass to determine rotational distance (Supplementary Figure 10). Rotational inertia was calculated, normalized to differences in humeral centroid size, and the inverse taken due to lower inertia values representing higher performance.

#### Humeral Torsion

The presence of humeral torsion - an angular offset between the proximal and distal articulations - has been hypothesized to increase effective stride length in sprawling taxa, by placing the manus further forward during walking, proportional to the sine of the torsion angle^31^. Given its association with sprawling locomotion, we predicted that humeral torsion would decrease throughout synapsid evolution^82^. We measured humeral torsion in 3-Matic by fitting two planes along the humerus – one aligned with the humeral head passing through the greater and lesser tubercles (or the homologous attachment points of the subscapularis and supracoracoideus muscles), and one aligned with the ulnar and radial condyles (Supplementary Figure 10). The angle between these two planes was then taken as the metric of humeral torsion and transformed by taking the sine of the angle.

#### Muscle Leverage

The anatomical and geometric arrangement of forelimb muscles are important determinants of functional performance. Muscle lever arms provide useful functional correlates for muscle action, as they directly correlate with muscle force and torque produced^83,84^. Of the muscle attachment sites easily identifiable across our humerus sample, the deltopectoral crest was the most reliable, and the deltoid and pectoral muscles that attach to it are both large muscles with important locomotor functions^14,37,41,74^. Therefore, we focused on the deltopectoral crest in our measurements of muscle leverage. Muscle in-levers were measured in 3-Matic from the most extreme point of the deltopectoral crest to the center of rotation at the humeral head (see the definition of rotational inertia), and then decomposed into separate components about different anatomical axes (Supplementary Figure 10).

For the X and Y axes, corresponding to humeral rotation in a vertical and horizontal plane, we calculated mechanical force advantages by dividing the muscle in-levers by humeral length (the out-lever for both these axes). We also measured velocity advantage by taking the inverse of force mechanical advantage (see note below). In both force and velocity advantages, the X and Y axis values were found to tightly correlate with one another, so these were averaged to produce single force and velocity advantage values. As humeral length is not the out-lever for long-axis rotation, and so cannot be used to derive mechanical advantage, in-levers for the corresponding Z axis, were size normalized by dividing by humerus centroid size. This left us with three size-normalized metrics of muscle leverage: ‘swing’ force leverage (combining the X and Y axis force advantages) to move the limb through a vertical or horizontal arc with high torque; ‘swing’ speed leverage (combining the X and Y axis velocity advantages) to move the limb through a vertical or horizontal arc with high speed; and ‘spin’ leverage (Z axis) to rotate the humerus about its long axis with high torque.

Adaptive landscape analyses often include traits that trade-off strongly with one another, but rarely are they exact inverses. While using separate surfaces for force and velocity may present methodological issues when few traits are models, it is less of a concern when multiple traits are model, as is the case for the present study. Further, it makes biological sense to include them both as individual traits which can be optimized as part of a combinatorial adaptive landscape, and we anticipated that some species would trade-off force for speed^14^.

#### Bending Strength

Limbs are subjected to varying loads during locomotion, and humeral strength determines the maximum external load that the bones can withstand. Although strength of the humerus is affected by both internal and external geometry^85,86^, it has been shown that external geometry alone can provide a good approximation of whole bone mechanical properties, especially across broad phylogenetic samples that vary widely in gross morphology^87^. Additionally, limitations of how fossil specimens could be digitized (surface scanning and photogrammetry rather than computed tomography) necessitated restricting the analysis to surface structures. Therefore, our analysis of bending strength only relies on external geometry. When meshes were imported into R, the modified *Morphomap* code calculated mechanically relevant cross-sectional properties of each landmarked slice, effectively assuming that the interior of the bone was entirely solid. Of these properties, we chose second moment of area at mid-shaft as our proxy for strength, as this is a commonly used metric for bending resistance in bones^86,88^. Second moment of area has units in dimensions of length to the fourth power, so to normalize for humerus size we took the fourth root of our second moment values before dividing by centroid size.

### Functional Traits and Performance Surfaces

Performance surfaces were created using the R package *Morphoscape*^21^. For each of the seven functional traits measured, values were standardized by scaling them to range between 0 and 1, with 0 representing lowest measured performance, and 1 representing highest measured performance. Functional traits measured on humerus models from all specimens were mapped onto the morphospace using ordinary Kriging to interpolate trait values across the space. This resulted in seven unique performance surfaces, which clearly show high-level performance gradients and form-function relationships (Figure 2) but are more ‘rugose’ than those from other studies^17,18,21^. Kriging as an interpolation method tends to produce more rugose surfaces than others (e.g., polynomial surface fitting), but it also makes no *a priori* assumptions about the number of performance peaks or the shape of the performance surface. It should also be noted that while the landmarking protocol is homology-free, the functional measurements (e.g., torsion and muscle leverage) are not, which may cause some occasional disconnects between form and function visible as local hills and valleys in the performance surface (see Supplementary Figure 11).

### Adaptive Landscapes

Adaptive landscapes were created using the R package *Morphoscape*. For each major taxonomic group of interest, landscapes were calculated by iteratively summing the seven performance surfaces, where each one had their contribution to overall performance weighted. All possible combinations of weights were tested, ranging from 0 to 1 in increments of 0.05, which for seven functional traits resulted in a total of 230,230 possible adaptive landscapes. Combinations of weights were favored that maximized the height of the point of interest – a group mean, an individual taxon or a reconstructed ancestral node - on the resulting landscape. For some taxa (e.g., reptiles and therians), despite clear functional differences between them, the top landscapes tended to have similar combinations of weights; therefore, all final landscapes presented here are based on the mean of the top 10% of weight combinations.

### Postural and Transitional Landscapes

To trace the evolution of the humerus as synapsids transitioned from sprawling to parasagittal limb postures, we first calculated an extant ‘sprawling’ vs ‘parasagittal’ landscape. As therians were the only group in our analysis with parasagittal posture, the parasagittal landscape is equivalent to the therian adaptive landscape. However, the sprawling groups differed significantly from each other, and were more broadly distributed across morphospace. Therefore, we created a composite sprawling landscape by overlaying the adaptive landscapes of individual sprawling groups (‘herptiles’, monotremes and non-synapsid fossils) and choosing the highest value in each grid cell as the value for the new landscape. High values on this new, composite landscape therefore indicated high functional performance for some form of sprawling, whether that be sprawling like a monotreme, reptile, salamander or other tetrapod^12,19,32,41^.

To calculate a transitional landscape and determine relative performance for sprawling or parasagittal postures in synapsids and other tetrapods, the difference between these ‘sprawling’ and ‘parasagittal’ landscapes was calculated by subtracting the values in each grid cell on the ‘sprawling’ landscape from the ‘parasagittal’ landscape. The resulting transitional landscape varies around 0, with negative values indicating increased performance on a sprawling landscape, and positive values indicating increased performance on the parasagittal therian landscape. To evaluate possible shifts in postural evolution across the ‘sprawling-parasagittal’ transition, tip and node performance values were extracted from the transitional landscape, and plotted onto the phylogeny using ggtree^89^. Following initial inspection of the tree, we then formally tested for differences in postural regime using the *SURFACE* package in R^90^ to fit OU models for different evolutionary regimes for posture (scores on the transitional landscape). As SURFACE can be prone to favoring overly complex models ^91^, we applied an AIC threshold of 4 during the model fitting process, so that more complex models with additional regimes had to provide a substantial improvement in model fit to be accepted over simpler models ^92^.

### Pareto Landscapes

To determine whether synapsids evolved along optimal evolutionary trajectories through morphospace, maintaining maximum overall performance even if the performance traits themselves might change, we constructed a series of Pareto landscapes^23,42,93^. The grid cells in morphospace, which represent humerus morphologies, can be considered as solutions to an optimization problem, and they experience trade-offs between different performance metrics and fitness on different adaptive landscapes. A subset of these solutions will be Pareto optimal, i.e., no other solution has better or equal performance in all metrics ^43^. This optimal subset of solutions is assigned rank 1, and then removed from the list of solutions before a second optimal subset (assigned rank 2) is determined. This process continues until all solutions have been ranked, thereby generating a Pareto ranking system.

Each Pareto landscape was calculated based on combining two adaptive landscapes reconstructed for major nodes in the Synapsid phylogeny – Synapsida, Therapsida, Cynodontia, Prozostrodontia and Theria. We ranked all locations in morphospace, based on their performance on the adaptive landscape reconstructed for a major synapsid node, and that for the next more derived major node (e.g., Synapsida and Therapsida, Therapsida and Cynodontia, etc.) toward which they might presumably be evolving. Optimality was ranked using a modified Goldberg ranking system^23,94^, first calculating the optimal ranking (RO) and then ranking again with the optimality of each metric reversed (suboptimal ranking, RS), using the R package *Rpref* ^95^. We then used the following equation to calculate Pareto optimality:

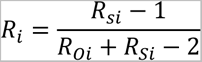

This equation produces a linear rank from 0 to 1, where 1 denotes Pareto optimal regions and 0 denotes the most suboptimal regions. We plotted the Pareto landscapes for four regions of the Synapsid phylogeny, based on differences in the adaptive landscapes reconstructed along the synapsid backbone (Figure 2C). We also mapped Pareto optimality calculated for each specific subset of non-mammalian synapsid taxa onto the phylogeny – tip values were taken from the Pareto landscape, and ancestral states were calculated via maximum likelihood.

## Acknowledgements

For assistance with specimen scanning and data acquisition we thank Jacqueline Lungmus, Spencer Hellert, Mark Wright, Peter Bishop, Blake Dickson, John Nyakatura, Tom Kemp, Roger Benson, and Elsa Panciroli. We thank Blake Dickson for assistance with *Morphoscape*. We are grateful to collections managers and curators from institutions all over the world who facilitated access to specimens - either physically or digitally - without whom this research would not have been possible. This work was funded by the NSF DEB 1754459 and NSF DEB 1754502, awarded to PIs S.E.P. and K.D.A., respectively. M.M. was supported by Grants-In-Aid of Undergraduate Research from the Harvard Museum of Comparative Zoology.

## Author Contributions

R.J.B. and S.E.P. conceived and designed the study. M.M. collected the anatomical and functional trait data, in consultation with R.J.B. and S.E.P., in partial fulfilment of an undergraduate senior thesis. R.J.B. wrote the R code and analyzed the morphological and functional data and drafted the figures. R.J.B. and S.E.P. interpreted the data and wrote the manuscript. All authors contributed to discussion, editing, and finalizing the text.

## Competing interests

The authors declare no competing interests.

## Notes

### Competing Interest Statement

The authors have declared no competing interest.

